# Hemolymph transfusions transfer heritable learned novel odor preferences to naïve larvae of *Bicyclus anynana* butterflies

**DOI:** 10.1101/2023.12.20.572719

**Authors:** V. Gowri, Antónia Monteiro

**Affiliations:** Department of Biological Sciences, National University of Singapore, 14 Science Drive 4 117543, Singapore

**Keywords:** *Bicyclus anynana*, Lepidoptera, odor learning, hemolymph, hemolymph transfer, inheritance of learned preference

## Abstract

The mechanisms whereby environmental experiences of parents are transmitted to their offspring to impact their behavior and fitness are poorly understood. Previously, we showed that naïve *Bicyclus anynana* butterfly larvae, whose parents fed on a normal plant feed but coated with a novel odor, inherited a learned preference towards that odor, which had initially elicited avoidance in the naïve parents. Here, we performed simple hemolymph transfusions from odor-fed and control-fed larvae to naïve larval recipients. We found that larvae injected with hemolymph from odor-fed donors stopped avoiding the novel odor, and their naïve offspring preferred the odor more, compared to the offspring of larvae injected with control hemolymph. These results indicate that factors in the hemolymph, potentially the odor molecule itself, play an important role in odor learning and preference transmission across generations. Furthermore, this mechanism of odor preference inheritance, mediated by the hemolymph, bypasses the peripheral odor-sensing mechanisms taking place in the antennae, mouthparts, or legs, and may mediate host plant switching and diversification in the Lepidoptera or more broadly across insects.

## 1. Introduction

Parental exposure to a novel stimulus or experience can influence the morphology, physiology, or behavior of their offspring. Such inheritance of acquired traits includes the development of a defensive taller helmet phenotype in water fleas [1,2], resistance to heavy metal toxicity in nematode worms [3], increased immunity against pathogens in *Galleria* moths [4], repulsive behavior towards a scent in pea aphids and mice [5,6], and preference for a novel odor or pheromone in fruit flies and butterflies [7-9]. All these studies show that the environment can influence an organism’s phenotype, and that of their offspring. However, the factors that are inherited are still illusive.

When organisms inherit a behavioral preference from their parents, this means that such a preference, expressed in the somatic cells of the central or peripheral nervous system of the parent, is transferred onto their germline (or accessory fluid cells) before being transmitted to their offspring. One vehicle for such transfer is the hemolymph, which circulates and is in direct contact with all the internal organs and tissues [10]. However, two possible behavioral preference inheritance mechanisms, mediated by hemolymph, should be considered. One involves the transfer of acquired preference factors from the brain to the germline and later to the offspring, to impact offspring behavior. The other involves the transfer of the same molecular stimulus that produced the behavioral change in the parent also to the germline, via retention of that stimulus in the hemolymph. While concrete evidence supporting either mechanism is lacking, hemolymph is the likely carrier of factors involved in both mechanisms. Thereby, transfusing it from treated to naïve individuals, and observing an altered phenotype in the latter, and/or in their offspring, can be a first step in isolating candidate factors that transfer a learned preference across individuals, and/or from parents to offspring.

Previous experiments with hemolymph transfers have shown that hemolymph can indeed transfer a ‘memory’ of a stimulus from experienced to naive recipients, altering the recipient’s phenotype [11-15]. But while these studies show that factors in the hemolymph can regulate immunity, behavior, wing color patterns, and longevity in the recipients of transfusions, there are barely any studies that explore the effects of hemolymph transfer across generations. It is thus, still unclear whether behavioral preferences acquired by a parental generation can be transferred to the offspring via hemolymph transfusions.

*Bicyclus anynana* is a subtropical model nymphalid butterfly that can learn preferences for novel visual cues [16,17], and transmit learned chemical cues to their offspring [8,18]. In previous studies, we showed that larvae that learned a novel food odor preference via feeding on that odor, produced naïve offspring that inherited that odor preference [8,19].

In this study, we investigate if the hemolymph of *B. anynana* larvae contains factors that can transfer learned odor preferences from odor-experienced to naïve larvae, and subsequently also to their offspring, via simple hemolymph transfusions. We fed larvae with odor-coated or control leaves throughout their larval stage. We then transfused hemolymph from each of these larval groups to naïve recipients. These recipients were tested for their odor preferences pre- and post-hemolymph transfusions. Once the recipient larvae became adults, they mated within each group, and their naïve larval offspring were tested for odor preferences. We hypothesized that factors contained in the hemolymph, either the odor itself or molecules produced in the brain downstream of odor feeding, can alter the odor preferences of both the recipient larvae and their offspring.

## 2. Material and methods

### (a) Animal husbandry

*B. anynana* were reared in a climate-controlled room at 27°C, 60% humidity, and 12:12-hour light: dark photoperiod. Butterflies were fed on mashed bananas. Corn leaves were placed in adult cages and wild-type embryos were collected from them after three to four hours.

### (b) Selecting larvae that showed a majority preference towards control or odor

Once larvae hatched from eggs (day 0), their naïve odor preferences were tested using odor choice assays (figure 1a; electronic supplementary material), and depending on the choice made, those larvae were assigned to either control or odor treatments. The odor used in this study is isoamyl acetate (IAA) which smells like bananas. Larvae of the control group were fed control-coated leaves and those of the odor group were fed odor-coated leaves (additional details in electronic supplementary material). Each larva was reared in a separate plastic container and tracked individually. The larvae were tested again for their odor choices on days 5, 10, 15 and 20. Only larvae that consistently chose the odor they were reared on (either control or odor), for at least four out of the five times they were tested, were selected as the ones that showed a ‘majority preference’ (figure 1b). These 20-day-old (5^th^ instar) larvae were used as donors in hemolymph transfusions.

**Figure 1.**
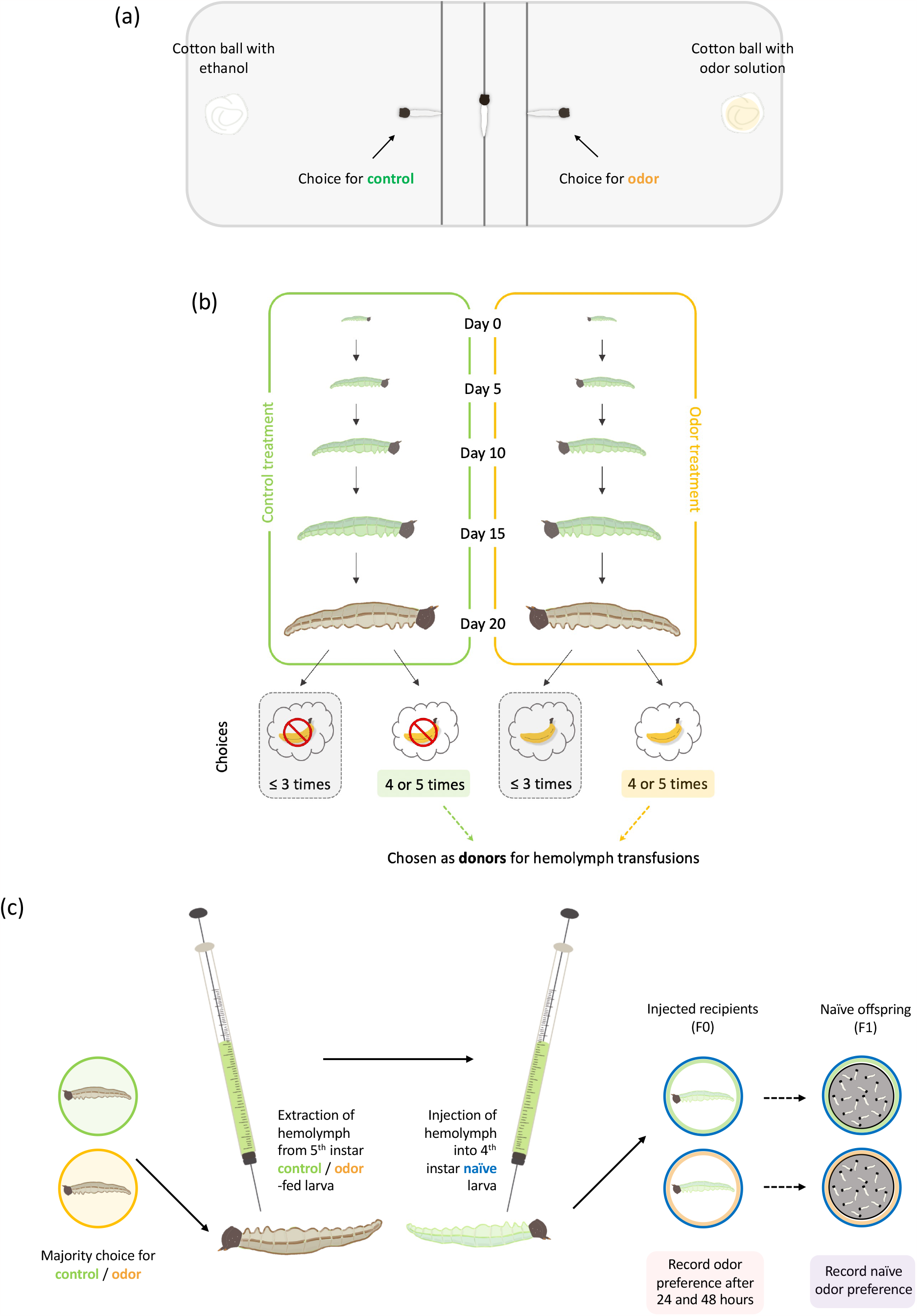
Experimental design. (a) *B. anynana* larvae were tested for either control or odor preference using the odor choice assay (additional details in electronic supplementary material). (b) Larvae that showed a consistent preference for the treatment odor they were reared on (at least 4 out of 5 choices throughout their development), were chosen as donors for hemolymph transfusions. (c) Hemolymph from control/odor treatment larvae was extracted and injected into naïve larvae. Odor preferences were recorded in the recipient larvae (24 and 48 hours after the transfusion) and in their naïve offspring.

### (c) Hemolymph transfusions and post-transfusion choice assays

Approximately 6μl of hemolymph was extracted from donor individuals and injected into recipient larvae (figure 1c). Injected recipient larvae were reared individually in plastic cups on uncoated leaves. Choice assays were performed at 24 hours and, again, at 48 hours after the hemolymph transfusion (HT). After pupation, the pupae were placed into two mesh group cages, depending on the injection treatment. Once the adult butterflies eclosed, they were allowed to mate within their treatment cage. Adults laid eggs on corn leaves placed in the cages, and eggs were then transferred onto Petri plates. As soon as these offspring larvae hatched, choice assays were performed to determine their naïve odor choices (figure 1c).

### (d) Statistical analyses

For all statistical analyses, only data from larvae that made a choice were used (electronic supplementary material, table S1).

#### Testing for larval odor preferences

To test if the proportion of larvae of each treatment choosing odor over control was significantly different from a random 50-50% choice, we used a chi-squared test of goodness of fit. Larvae were considered to have a ‘preference’ if their choice deviated significantly from 50%.

#### Testing for differences in odor choice made by recipient larvae over time and between treatments

We tested the effect of hemolymph type (from control or odor-fed donors) and the time point of odor choice assay (24- and 48-hours post-transfusion) on larval odor choice, by conducting repeated measures ANOVA on binomial generalized linear mixed-effects model (GLMER). For this specific test, only data from larvae that survived and made a choice till 48 hours post HT was used. In the logit link function, we coded the choice for control as 0, and the choice for odor as 1. We tested for factors that contributed to explaining variation in the response variable using likelihood ratio tests (LRT). Subsequently, we removed the non-significant factor from the final model. The differences in choices made by the larvae pre- and post-HT of each treatment were compared using two-tailed Fisher exact test of independence.

#### Testing the effect of hemolymph transfusion on offspring larvae

The difference in the choices made by the naïve offspring larvae between treatments were compared using two-tailed Fisher exact test of independence.

All analyses were performed in the R statistical framework (RStudio Team 2022), using the packages lsmeans [20], lme4 [21], rcompanion [22], car [23], multcompView [24], and Rmisc [25].

## 3. Results

### Larvae injected with hemolymph from odor-fed donor larvae showed a significant change in their odor preference, and so did their offspring

Before hemolymph transfusions (HT), naïve larvae showed a significant preference for control relative to odor (37% chose odor; *n*_naïve_ = 112, chi-squared = 8.036, df = 1, *P*-value = 0.005; figure 2). 24 and 48 hours after HT, control (C_24_ or C_48_) and banana odor-recipient larvae (B_24_ or B_48_) made random choices, i.e., they showed neither a preference for control nor for odor, when tested at each time point (45% of C_24_ chose odor: *n* = 33, chi-squared = 0.273, df = 1, *P*-value = 0.602; 64% of B_24_ chose odor: *n* = 39, chi-squared = 3.103, df = 1, *P*-value = 0.078; 35% of C_48_ chose odor: *n* = 20, chi-squared = 1.8, df = 1, *P*-value = 0.180; 61% of B_48_ chose odor: *n* = 28, chi-squared = 1.286, df = 1, *P*-value = 0.257; figure 2).

**Figure 2.**
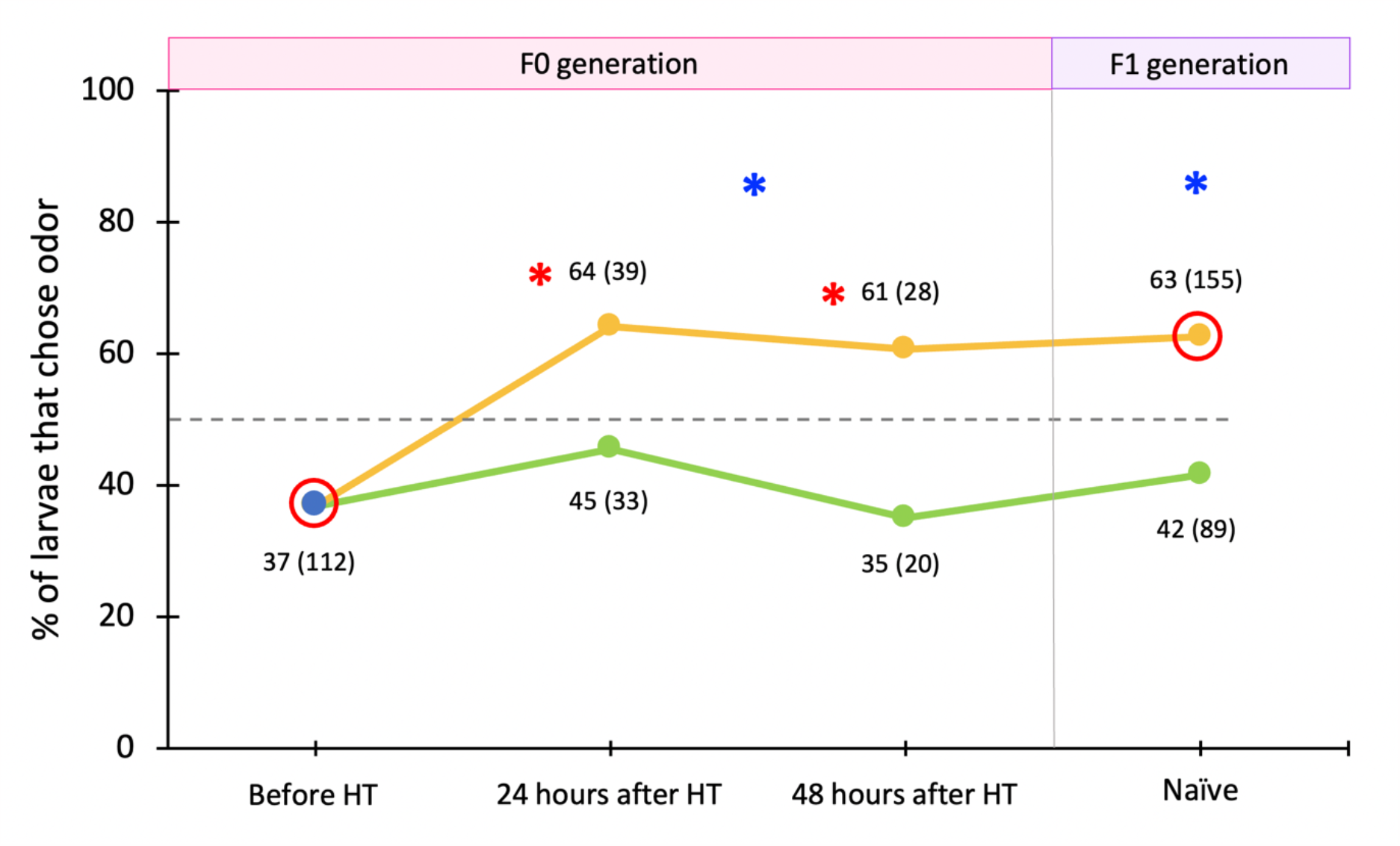
Odor preferences of *B. anynana* larvae before and after hemolymph transfusions (HT). Naïve recipient larvae (blue dot), with an initial significant avoidance of the novel odor, were later monitored for odor preferences post-transfusion. Green and orange lines represent larvae injected with hemolymph from either control or odor-fed larvae, respectively. Near each point, the percentage values of choice for odor are denoted along with the corresponding total sample sizes (larvae that made a choice) in brackets. Significant preferences (deviation from a random choice) are represented by red circles that outline those specific data points. Red asterisks denote a significantly different choice relative to that made by the larvae before HT. Blue asterisks denote that the choices of both injected recipient and offspring groups are significantly different between treatments (*, *P*-value<0.05).

B_24_ and B_48_ larvae showed significantly increased choices for odor relative to pre-injected larvae (Fisher exact test, 24 hours after HT: *n*_naïve_ = 112, *n*_B_ = 39, *P*-value = 0.005; 48 hours after HT: *n*_naïve_ = 112, *n*_B_ = 28, *P*-value = 0.031; figure 2). C_24_ and C_48_ larvae, however, did not show such shifts relative to pre-injected larvae (Fisher exact test, 24 hours after HT: *n*_naïve_ = 112, *n*_C_ = 33, *P*-value = 0.418; 48 hours after HT: *n*_naïve_ = 112, *n*_C_ = 20, *P*-value = 1; figure 2). Odor hemolymph-injected larvae chose the odor significantly more compared to control hemolymph-injected larvae (post hoc comparison, B-C: *p* = 0.0169; figure 2, Table 1).

**Table 1:** Summary of repeated measures ANOVA on GLMER (figure 2).

| | Chisq | df | $P$-value |
| --- | --- | --- | --- |
| **Odor choice of recipient larvae** |  |  |  |
| **Full model variables** |  |  |  |
| Treatment (hemolymph type C or B) | 0.076 | 1 | 0.783 |
| Time point (24- or 48-hrs post transfusion) | 2.711 | 1 | 0.1 |
| Treatment x time | 0.005 | 1 | 0.943 |
| **Final model variable** |  |  |  |
| Treatment | 5.708 | 1 | 0.017 |

Offspring of parents injected with hemolymph from odor-fed larvae (B_off_), however, showed a significant preference for odor (63% of B_off_ chose odor: *n* = 155, chi-squared = 9.813, df = 1, *P*-value = 0.002; figure 2), whereas offspring of larvae injected with hemolymph from control-fed larvae (C_off_) made a random choice (42% of C_off_ chose odor: *n* = 89, chi-squared = 2.528, df = 1, *P*-value = 0.118; figure 2). In addition, B_off_ larvae preferred odor significantly more than the C_off_ larvae (Fisher exact test, naïve offspring: *n*_C_ = 89, *n*_B_ = 155, *P*-value = 0.002; figure 2).

## 4. Discussion

In this study, we performed hemolymph transfusion experiments from larvae fed on different diets, one of them containing a novel odor, to naïve *B. anynana* larvae to investigate if the hemolymph can influence the odor preferences of recipients and those of their offspring. We showed that the hemolymph of odor-fed larvae contains factors that can change the behavior of recipient larvae as well as that of their offspring: it makes them move toward that odor. This study shows the transfer of a learned odor preference across lepidopteran individuals and generations using hemolymph transfusions.

Changes in phenotype have been previously documented after hemolymph transfusion studies [11-15], but our study, documenting a behavioral change, mostly resembles a previous hemolymph transfusion study done in mosquitoes [13]. Here, hemolymph from egg-carrying female mosquitos led non-egg-carrying females to halt their host-seeking behavior [13]. The factors in the hemolymph in these egg-carrying female mosquitoes, likely by-products of internal changes in physiology, got transferred to host females to affect their behavior. In contrast, our study shows that hemolymph transfusions created preferences towards external cues, such as odors, completely bypassing stereotypical odor sensory mechanisms in insects. These mechanisms typically involve the odor molecule binding to gustatory or olfactory receptors in the mouth parts or antennae, and the odor stimulus being transported from the peripheral nervous system to the brain. In addition, our experiment also showed that these odor learning-inducing factors in the hemolymph can also travel throughout the larval body, reach the germline, and impact odor preferences in the next generation of odor-naïve larvae.

It is still unclear, however, how the encounter with a novel odor induces heritable larval hemolymph chemistry changes. One possibility is the inheritance of the odorant molecules themselves. Odorants from the feed leaves might have directly entered the hemolymph via small openings on the larval cuticle/exoskeleton during respiration, sensory sensilla, or gut. Once dissolved in the hemolymph, odor molecules could have been transported to the germ cells, to later influence odor preferences in the offspring that developed from those cells. This hemolymph-mediated mechanism of odor preference development is supported by a study on pigeons where an increased olfactory response was observed upon injection of just the odorant into the blood of recipients [26].

Alternatively, the novel odorant might have induced heritable epigenetic changes in the hemolymph [27-29]. A possible mechanism can start with hemolymph-borne odorants binding to odorant-binding proteins (OBPs) in the antennae, and then being transported to the signal processing center of the brain by the circulating hemolymph. These protein-bound odorants might induce DNA methylation marks, histone acetylation, or expression of non-coding RNAs in the brain that then travel back to the hemolymph and the germ line. Dias and Ressler suggested that blood-borne odorants might activate olfactory receptors in the germ cells directly, and proposed this as a possible mechanism for the inheritance of odor aversion observed in mice [6]. Other mechanisms might include the inheritance of epigenetic factors such as methylation marks on top of specific odor receptors and microRNAs (miRNAs) upon odor-fear conditioning in mice [6,30], starvation-induced small RNAs in nematode worms [31], and piwi-interacting RNAs (piRNAs) that aid in pathogen-avoiding behavior in worms [32].

Given the diversity of possible mechanisms connected to odor preference/avoidance inheritance, future studies may need to i) extract, isolate and transfuse different factors from the hemolymph in isolation into naïve individuals to observe whether they are responsible for the development of an odor preference, and ii) explore the epigenetic factors involved in odor learning and inheritance of learned odor preferences. Future studies with *B. anynana* larvae can also be improved by making the choice arena Y-shaped and provided with continuous airflow, to accentuate the difference between odor choices.

## Ethics

This work did not require ethical approval from a human subject or animal welfare committee.

## Data accessibility

The data associated with this study is available in the Supplementary material.

## Declaration of AI use

We have not used AI-assisted technologies in creating this article.

## Author contributions

VG and AM designed the study. VG performed the experiments, collected and analyzed all the data, and made the figures. VG and AM wrote the manuscript and contributed to revisions.

## Acknowledgements

We thank the Fire Flies Health Farm and Greenology for providing corn plants. We are grateful for Emilie Dion and Ajay Sriram Mathuru for their valuable support and suggestions regarding the statistical analysis of this work.

## References

1. Tollrian, R. (1990). Predator-induced helmet formation in Daphnia cucullata (Sars). Archiv Für Hydrobiologie, 119(2), 191–196.

2. Tollrian, R. (1995). Chaoborus crystallinus predation on Daphnia pulex: can induced morphological changes balance effects of body size on vulnerability? Oecologia, 101(2), 151–155.

3. Kishimoto, S., Uno, M., Okabe, E., Nono, M., & Nishida, E. (2017). Environmental stresses induce transgenerationally inheritable survival advantages via germline-to-soma communication in Caenorhabditis elegans. Nature Communications, 8(1), 14031.

4. Freitak, D., Schmidtberg, H., Dickel, F., Lochnit, G., Vogel, H., & Vilcinskas, A. (2014). The maternal transfer of bacteria can mediate trans-generational immune priming in insects. Virulence, 5(4), 547–554.

5. Keiser, C. N., & Mondor, E. B. (2013). Transgenerational Behavioral Plasticity in a Parthenogenetic Insect in Response to Increased Predation Risk. Journal of Insect Behavior, 26(4), 603–613.

6. Dias, B. G., & Ressler, K. J. (2014). Parental olfactory experience influences behavior and neural structure in subsequent generations. Nature Neuroscience, 17(1), 89–96.

7. Williams, Z. M. (2016). Transgenerational influence of sensorimotor training on offspring behavior and its neural basis in Drosophila. Neurobiology of Learning and Memory, 131, 166–175.

8. Gowri, V., Dion, E., Viswanath, A., Piel, F. M., & Monteiro, A. (2019). Transgenerational inheritance of learned preferences for novel host plant odors in Bicyclus anynana butterflies. Evolution, 73(12), 2401–2414.

9. Dion, E., Pui, L. X., Weber, K., & Monteiro, A. (2020). Early exposure to new sex pheromone blends alters mate preference in female butterflies and their offspring. Nature Communications, 11(1), 53.

10. Hillyer, J. F., & Pass, G. (2020). The insect circulatory system: structure, function, and evolution. Annual Review of Entomology, 65, 121–143.

11. Rodrigues, J., Brayner, F. A., Alves, L. C., Dixit, R., & Barillas-Mury, C. (2010). Hemocyte differentiation mediates innate immune memory in Anopheles gambiae mosquitoes. Science (New York, N.Y.), 329(5997), 1353–1355.

12. Wiesner, A. (1991). Induction of immunity by latex beads and by hemolymph transfer in Galleria mellonella. Developmental & Comparative Immunology, 15(4), 241–250.

13. Klowden, M. J., & Lea, A. O. (1979). Humoral inhibition of host-seeking in Aedes aegypti during oöcyte maturation. Journal of Insect Physiology, 25(3), 231–235.

14. Otaki, J. M. (1998). Color-pattern modifications of butterfly wings induced by transfusion and oxyanions. Journal of Insect Physiology, 44(12), 1181–1190.

15. Sondhi, K. C. (1967). Studies in aging, 3. The physiological effects of injecting hemolymph from outbred donors into inbred hosts in Drosophila melanogaster. Proceedings of the National Academy of Sciences, 57(4), 965–971.

16. Westerman, E. L., Hodgins-Davis, A., Dinwiddie, A., & Monteiro, A. (2012). Biased learning affects mate choice in a butterfly. Proceedings of the National Academy of Sciences, 109(27), 10948 LP –10953.

17. Westerman, E. L., & Monteiro, A. (2013). Odour influences whether females learn to prefer or to avoid wing patterns of male butterflies. Animal Behaviour, 86(6), 1139–1145.

18. Dion, E., Pui, L. X., & Monteiro, A. (2017). Early-exposure to new sex pheromone blend alters mate preference in female butterflies and in their offspring. bioRxiv, 214635.

19. Gowri, V., & Monteiro, A. (2023). Acquired preferences for a novel food odor do not become stronger or stable after multiple generations of odor-feeding in Bicyclus anynana butterfly larvae. Annals of the New York Academy of Sciences, 00, 1–11.

20. Lenth, R., & Lenth, M. R. (2018). Package ‘lsmeans.’ The American Statistician, 34(4), 216–221.

21. Bates, D., Kliegl, R., Vasishth, S., & Baayen, H. (2015). Parsimonious mixed models. ArXiv Preprint ArXiv:1506.04967.

22. Mangiafico, S. S. (2016). Summary and Analysis of Extension. Program Evaluation in R, Version, 1(1).

23. Fox, J., & Weisberg, S. (2009). car: companion to applied regression. R package version 1: 2–14. Available at http.Cran.Rproject-Org/Web/Packages/Car/Index.Html.

24. Graves, S., H.-P. Piepho, and L. Selzer. 2015. multcompView: visualizations of paired comparisons. R package version 0.1–7 (with help from Sundar Dorai-Raj). Available at https://CRAN.R-project.org/package=multcompView.

25. Hope, R. M., Hope, M. R. M., & Collate’CI, R. (2013). Package ‘Rmisc.’ Group, 101, 2.

26. Maruniak, J. A., Silver, W. L., & Moulton, D. G. (1983). Olfactory receptors respond to blood-borne odorants. Brain Research, 265(2), 312–316.

27. Danchin, E., Pocheville, A., Rey, O., Pujol, B., & Blanchet, S. (2019). Epigenetically facilitated mutational assimilation: epigenetics as a hub within the inclusive evolutionary synthesis. Biological Reviews of the Cambridge Philosophical Society, 94(1), 259–282.

28. Gowri, V., & Monteiro, A. (2021). Inheritance of Acquired Traits in Insects and Other Animals and the Epigenetic Mechanisms that Break the Weismann Barrier. In Journal of Developmental Biology (Vol. 9, Issue 4).

29. Villagra, C., & Frías-Lasserre, D. (2020). Epigenetic molecular mechanisms in insects. Neotropical Entomology, 49(5), 615–642.

30. Aoued, H. S., Sannigrahi, S., Hunter, S. C., Doshi, N., Sathi, Z. S., Chan, A. W. S., Walum, H., & Dias, B. G. (2020). Proximate causes and consequences of intergenerational influences of salient sensory experience. Genes, Brain and Behavior, 19(4), e12638.

31. Rechavi, O., Houri-Ze’evi, L., Anava, S., Goh, W. S. S., Kerk, S. Y., Hannon, G. J., & Hobert, O. (2014). Starvation-Induced Transgenerational Inheritance of Small RNAs in C. elegans. Cell, 158(2), 277–287.

32. Kaletsky, R., Moore, R. S., Vrla, G. D., Parsons, L. R., Gitai, Z., & Murphy, C. T. (2020). C. elegans interprets bacterial non-coding RNAs to learn pathogenic avoidance. Nature, 586(7829), 445–451.

