## Supplementary material for "Hemolymph transfusions transfer heritable learned novel odor preferences to naïve larvae of *Bicyclus anynana* butterflies": Supplementary information_Gowri&Monteiro.docx

**Odor choice assay**

An odor choice assay was used to determine the odor preferences of larvae (figure 1a). The choice assay arena consisted of a white plastic board, with a line drawn in the middle. A line was drawn on either side of the midline, at a distance corresponding to the length of the larval body at the time of the choice assay. Five drops of the control solution were added to a cotton ball that was placed at one end, 12cm away from the midline. Similarly, five drops of the odor solution were added to another cotton ball and placed at the other end. Each larva was aligned along the midline, and a white translucent plastic box, measuring 26.5cm X 9cm X 5cm was placed over the arena to cover and prevent any interfering visual and odor cues. A small piece of green tape was stuck at each (inner) end of the box so that the larvae, who are attracted to that color, move towards the ends of the arena and not towards the sides. Larvae at days 0, 5 and 10 were given four minutes to make a choice. Larger larvae, on days 15 and 20, were given five minutes to make a choice. At the end of these time intervals, if the larvae had crossed the either line, this event was determined a ‘choice for control’ or a ‘choice for odor’, depending on the side it walked towards. If no choice was made at the end of the time interval, this was noted as ‘no choice’.

**Preparation of scented leaves**

Banana-smelling organic compound isoamyl acetate (IAA), which is also known as isopentyl acetate, from Sigma-Aldrich (W205532, natural, ≥97%, Food Chemicals Codex, Food Grade) was used as the novel odor in this study. Since IAA is not highly soluble in water, absolute ethanol (Fisher Chemical 99.8%, Analytical reagent grade) was used as the solvent in odor solutions preparations. 5% IAA was used as the odor solution, and absolute ethanol was used as the control solution. Cotton saturated with the solutions was used to rub and coat the cut corn leaves thoroughly. Once the ethanol completely evaporated, the leaves were used to feed the larvae. Control solution-coated leaves (control leaves) and odor solution-coated leaves (odor leaves) were used as the larval feed for the control treatment and odor treatment respectively, till pupation. To make sure that the larvae were continuously exposed to the odor, the feed was replaced once every two days with leaves that were freshly coated with the solutions.

**Hemolymph transfusion experiment**

Fourth instar naïve larvae from the common lab stock were chosen as recipient larvae. These larvae are big enough to take in a considerable volume of hemolymph, yet young enough to have time to recover post-injection, and still be tested for a change in preference before they pupate. We injected only approximately 6µl of hemolymph as we found via trials that a higher volume led to complications in larval development and increased mortality. Innate odor choice for each recipient larva was determined using a single choice assay, prior to hemolymph injection. For the extraction and injection of hemolymph, Hamilton needles (gauge 33, point 4, 10mm, angle 12°) and Hamilton syringes (Gastight, 1700 series, 10µl) were used.

Each donor/recipient larva was kept on ice for no more than one minute to immobilize them slightly. A small puncture was made on the mid-section of the donor larval body using a disposable needle and the body was squeezed with slight pressure to release some hemolymph. Approximately 6µl of hemolymph was extracted from the donor individual and injected into the recipient larva (figure 1c). This was done immediately to avoid hemolymph degradation and melanization. Slight pressure was applied to the site of injection of the recipient larva using a finger, for a few seconds, to allow for some healing and to hold the fluid inside the body before the larva was put back in its container. In some instances, the same donor larva was used for up to three different transfusions, in short succession, for a maximum of 18ul of hemolymph drawn.

**Supplemental Table S1. Odor choices made by recipient larvae before and after hemolymph transfusions (HT), and by their naïve offspring of both treatments.**

| **Donor treatment** | **Time point of choice assay** | **Choice for control** | | **Choice for odor** | | **Total #** |
| --- | --- | --- | --- | --- | --- | --- |
|  |  | **No.** | **%** | **No.** | **%** | **No.** |
| Control | Before HT | 31 | 61 | 20 | 39 | 51 |
|  | 24 hours after HT | 18 | 55 | 15 | 45 | 33 |
|  | 48 hours after HT | 13 | 65 | 7 | 35 | 20 |
|  | Naïve offspring | 52 | 58 | 37 | 42 | 89 |
| Odor | Before HT | 40 | 66 | 21 | 34 | 61 |
|  | 24 hours after HT | 14 | 36 | 25 | 64 | 39 |
|  | 48 hours after HT | 11 | 39 | 17 | 61 | 28 |
|  | Naïve offspring | 58 | 37 | 97 | 63 | 155 |
